# Privacy-preserving genotype imputation with fully homomorphic encryption

**DOI:** 10.1101/2020.05.29.124412

**Authors:** Gamze Gürsoy, Eduardo Chielle, Charlotte M. Brannon, Michail Maniatakos, Mark Gerstein

**Affiliations:** Program in Computational Biology and Bioinformatics, Yale University, New Haven, CT 06520, USA; Department of Molecular Biophysics and Biochemistry, Yale University, New Haven, CT 06520, USA; Department of Electrical and Computer Engineering, New York University Abu Dhabi, Abu Dhabi, UAE; Department of Computer Science, Yale University, New Haven, CT 06520, USA

**Keywords:** genome privacy, genotype imputation, homomorphic encryption

## Abstract

Genotype imputation is the statistical inference of unknown genotypes using known population haplotype structures observed in large genomic datasets, such as HapMap and 1000 genomes project. Genotype imputation can help further our understanding of the relationships between genotypes and traits, and is extremely useful for analyses such as genome-wide association studies and expression quantitative loci inference. Increasing the number of genotyped genomes will increase the statistical power for inferring genotype-phenotype relationships, but the amount of data required and the compute-intense nature of the genotype imputation problem overwhelms servers. Hence, many institutions are moving towards outsourcing cloud services to scale up research in a cost effective manner. This raises privacy concerns, which we propose to address via homomorphic encryption. Homomorphic encryption is a type of encryption that allows data analysis on ciphertexts, and would thereby avoid the decryption of private genotypes in the cloud. Here we develop an efficient, privacy-preserving genotype imputation algorithm, p-Impute, using homomorphic encryption. Our results showed that the performance of p-Impute is equivalent to the state-of-the-art plaintext solutions, achieving up to 99% micro area under curve score, and requiring a scalable amount of memory and computational time.

## 1 Introduction

The decreasing cost of DNA sequencing technologies and the clinical importance of genomic characterization of individuals have resulted in an exponential increase of available human genetic data [1]. Such data collection has tremendous clinical value in terms of understanding and characterizing rare diseases and genotype-phenotype associations [2, 3]. However, the large amounts of data being collected and the complexity of genomic analyses overwhelm the capacity of servers. Therefore, funding agencies and institutions have begun outsourcing cloud services. For example, The National Human Research Institute Genomic Data Science Analysis, Visualization, and Informatics Lab-space (AnVIL) [4] provides a scalable cloud-based infrastructure for genomic data access, sharing and computing across large genomic, and genomic-related data sets. These cloud-based services raise privacy issues when used for sharing patients’ genomic data. With the increasing use of genetic information in new avenues, such as forensics, the need for privacy-preserving analysis methods is greater than ever.

Genotype imputation is the process of statistical inference of unknown genotypes in a genome using the correlation between the single nucleotide polymorphisms (SNP) sites observed in population-based haplotype structures. Genotype imputation has immense value in phenotype-genotype association studies, especially when the number of study participants is large. It allows researchers to perform cost-effective genotyping methods, such as genotyping arrays or low-coverage whole genome sequencing, and obtain missing or low-quality genotypes through statistical inference. As the input and output of genotype imputation methods are sets of participants’ genotypes, performing this analysis in the cloud could have serious privacy implications, such as exposure of the sensitive data to malicious parties. On the other hand, the large amounts of data required to boost statistical power makes local storage and analysis cost prohibitive. Therefore, it is essential to develop privacy-preserving genotype imputation methods that allow users to utilize cloud-based services while protecting privacy. Here we develop a privacy-preserving genotype imputation method called *p-Impute* based on homomorphic encryption (HE). p-Impute allows users to perform genotype imputation on encrypted genotype data and returns encrypted genotype outputs, hence removing the privacy concerns related to outsourcing cloud services.

Homomorphic encryption (HE) is a form of encryption with the capability for computing on encrypted data without access to the secret key [5]. Homomorphic encryption schemes are classified based on the different kinds of computation that they can successfully perform on encrypted data. Partially homomorphic encryption (PHE) supports the evaluation of unbound arithmetic circuits consisting of only one type of operation such as addition or multiplication only. Somewhat homomorphic encryption (SHE) supports the evaluation of two orthogonal types of operations, such as addition and multiplication, for circuits with known depth (i.e with known number of operations multiplication and/or addition in the algorithm). This limitation is due to the noise introduced to the ciphertexts, where each additional operation increases the noise in the resulting ciphertext. Fully homomorphic encryption (FHE) supports two orthogonal operations, such as addition and multiplication, and due to bootstrapping, has unbound computation. Bootstrapping is an expensive operation that reduces the noise of ciphertexts. For both SHE and FHE, it is important to optimize the circuit for depth, since the noise accumulated is related to the depth of the circuit. Additionally, addition operations are fast and produce little noise, while multiplications are slow and produce more noise. In summary, PHE is fast and unbound, but less expressive; SHE slow and bounded, but more expressive; and FHE is expressive and unbound, but very slow. Furthermore, in HE, control flow cannot depend on the data. For example, we cannot perform an *if* clause on encrypted data. Therefore, the program must run obliviously to its sensitive data, which leads to further performance degradation [6]. Although HE has been used for the analysis of genomic and biomedical clinical data before [7, 8, 9, 10], it requires a fast and scalable implementation to be used for genotype imputation.

In this study, we design *p-Impute*, a privacy-preserving statistical inference algorithm to perform genotype imputation using the BFV (Brakerski/Fan-Vercauteren) encryption scheme [11] provided by Microsoft SEAL [12] library. We overcome the challenges related to performance overhead through algorithm optimization, batching, and thread-level parallelism. We used 2,504 fully characterized genomes from 1000 genomes project [13] as training and predicted the genotypes for the target SNPs of 870 individuals from Genotype Tissue Expression project (GTEx) [14] as test. Moreover, we compared *p-Impute* results to the results from state-of-the-art non-encrypted counterparts (IMPUTE2 [15] and Beagle [16]) and found an excellent agreement, where all three algorithms resulted in 99% AUC on GTEx data.

## 2 Results

### 2.1 Scenario

Consider a scenario where Alice has a set of genotyped tag SNPs and would like to impute the genotypes of her missing SNPs (target SNPs) using the tag SNPs. She would like to outsource the imputation to Bob, as Bob has a genotype imputation model, but Alice does not trust Bob enough to send him her SNPs. Instead, Bob offers an imputation method on the encrypted space. Alice encrypts her tag SNP genotypes with her public key and sends to Bob. Bob runs his imputation method and gets encrypted results and sends them to Alice. Alice uses her private key to decrypt the results, which are her missing SNP genotypes (Figure 1). In a real-world scenario, Bob could be the cloud-service provider, who will never have access to any SNP from any individual.

**Figure 1:**
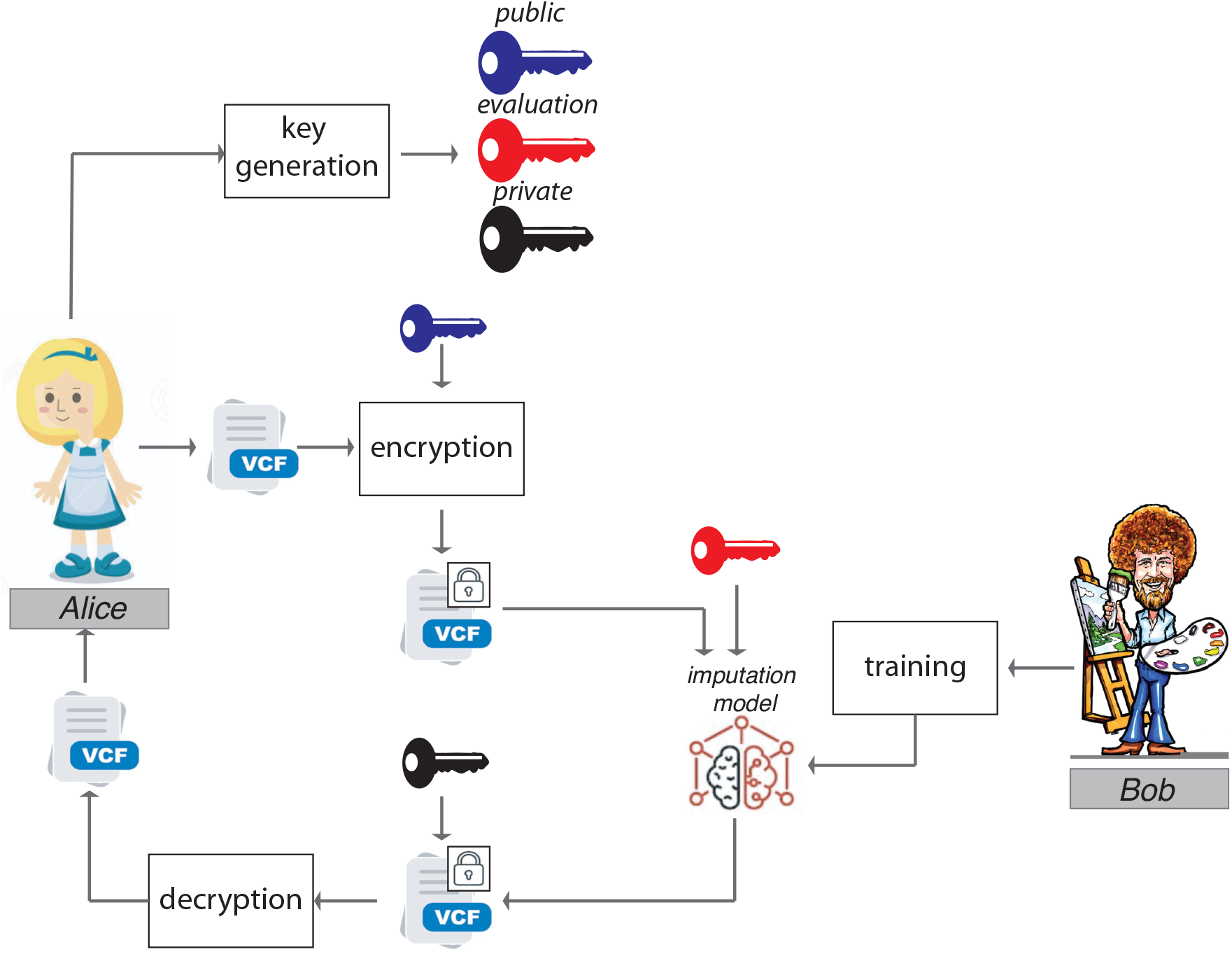
Threat Model. Illustration of how genotype imputation with homomorphic encryption works in practice. After generating encryption, decryption and evaluation keys, Alice encrypts the vcf file with her tag SNP genotypes with her encryption key and sends the encrypted genotypes and the evaluation key to Bob. Bob uses the encrypted genotypes as input to his privacy-preserving imputation model and outputs encrypted genotypes for Alice’s missing SNPs and sends them to Alice. Alice can decrypt the output using her decryption key. Bob never receives any unencrypted data and never sees the plaintext results from his model.

### 2.2 Dataset

We used fully characterized genomes from 2504 individuals provided by the 1000 Genomes Project [13] as our training dataset. Chromosome 1 of the human genome was divided into sets of tag and target SNPs by the iDASH Secure Genome Analysis Challenge’19 [17], which we used in this study. In total, for Chromosome 1, we had 9745 tag SNPs and 500 target SNPs. For the test dataset, we used the Chromosome 1 genotypes characterized by whole genome sequencing of 870 individuals in the GTEx project [14]. The task is to predict the genotypes of 500 target SNPs by using the genotypes of 9745 tag SNPs. We performed the same predictions by using only 1045 tag SNP genotypes for inferring 500 target SNP genotypes in order to see the effect of the number of tag SNPs on overall predictions. Both 9745 and 1045 tag SNPs were selected by iDASH Secure Genome Analysis Challenge’19 using different genomic distance cut-off between SNPs, i.e 9745 tag SNPs are 1kb apart from each other, whereas 1045 tag SNPs are 10kb apart from each other.

### 2.3 Genotype imputation algorithm design

Traditional plaintext genotype imputation methods such as IMPUTE2 [15] and Beagle [16] first phase the genome into haplotypes. They next determine the best haplotype block structure using a genotype panel from different populations. Haplotype blocks are then used to determine the relationship between tag and missing SNP genotypes in order to compute a probability for having each genotype (0, 1, and 2) at the missing SNP position. Many of the imputation methods available are based on Hidden Markov Models [15, 16, 18]. Complex statistical methods such as Hidden Markov Models require operations such as exponentiation, activation, or feedback. Thus, their implementation using HE-based principles for computation in the encrypted domain may not be able to scale-up genotype imputation for hundreds of individuals due to the large performance overheads.

To avoid some of the overhead problems, we develop a plaintext method that does not depend on phasing and can impute genotypes directly using the tag SNP genotypes (0,1 or 2). We describe our algorithm in Figure 2. For each missing target SNP genotype of a query individual, we determine *N* number of tag SNPs that are closest to it. We call this group of SNPs “pseudo-haplotype block”. We define a similarity score between the query individual and each individual *k* in the training database for the pseudo-haplotype block as,

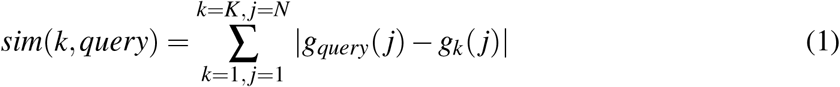

where *K* is the total number of individuals in the training database, *N* is the total number of tag SNPs in the pseudo-haplotype block, *g*_*query*_(*j*) is the genotype of the *j*^*th*^ tag SNP in the pseudo-haplotype block of the query individual, and *g*_*k*_(*j*) is the genotype of the *j*^*th*^ tag SNP in the pseudo-haplotype block of the *k*^*th*^ individual in the training database. After the calculation of the similarity between all the individuals and the query individual, we sort *sim*(*k, query*) scores in increasing order and take the top *M* individuals in the database as our training individuals. These individuals are genetically similar to the query individual within a pseudo-haplotype block. We then calculate a score for each genotype at the missing SNP location by calculating the joint probabilities of genotypes between the missing and tag SNPs as following.

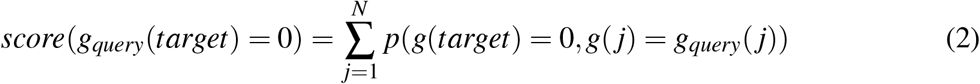

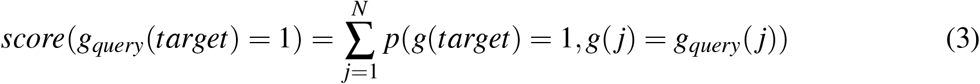

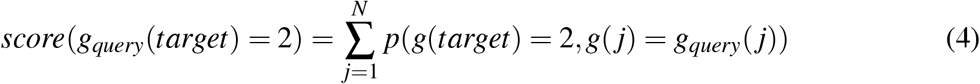

**Figure 2:**
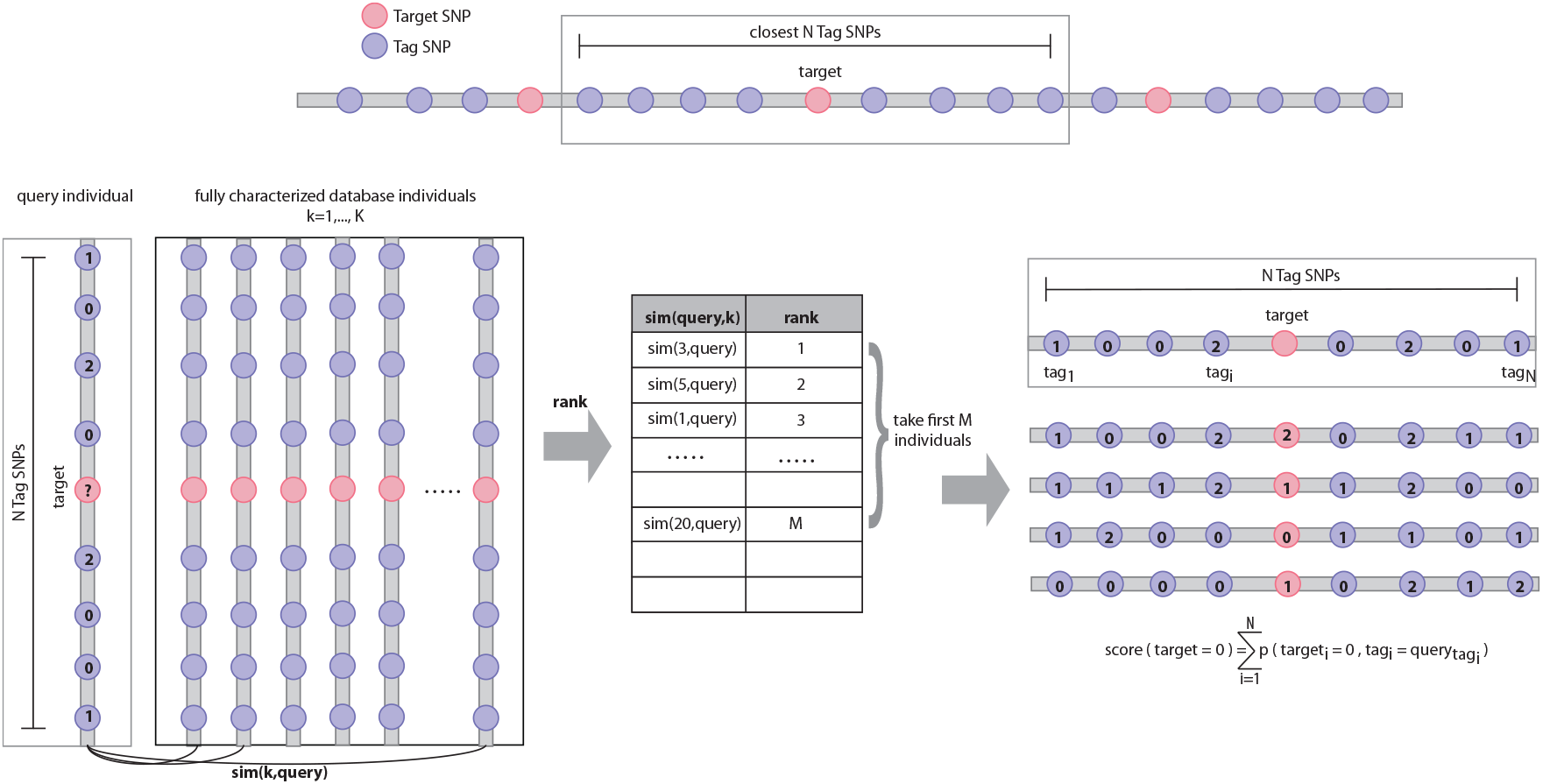
p-Impute algorithm. After determining the neighboring tag SNPs for the missing genotype, the similarity between the query individual and the each individual in the training database is calculated. The training individuals are ranked based on the similarity to the query individual and the top *M* individual are used to calculate the probability of each genotype being observed at the missing SNP location based on the probability of joint occurrence between the missing SNP location genotype and tag SNP genotypes in the training database.

We then normalize these scores to obtain probability of having a genotype at the missing SNP location. The probability of having the genotype *x* such that *x* = {0, 1, 2} at the missing SNP location becomes

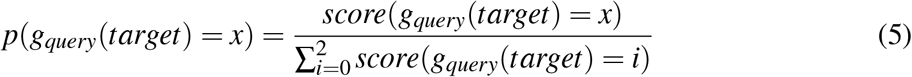

### 2.4 Tuning the algorithm for HE

#### Training

Here we update the algorithm described in Section 2.3 such that training does not require any private input, i.e. the genotypes of the query individual. We calculate the genotype probabilities for each target SNP of a query individual using the correlation of a group of selected tag SNPs between the query individual and database individuals. We select these tag SNPs by their proximity to the target SNP. Thus, we first must define the number *N* of tag SNPs we want to select. Since there are 3 possible values for the genotype (0, 1, or 2), there are 3^*N*^ possible combinations of tag SNPs. For each combination, we create a *virtual individual* containing those tag SNPs and then we apply the method described in Section 2.3. Once we calculate the most related database individuals (to the virtual query individual) using Eq. 1, we then use a subset of database individuals (the most similar ones) to calculate the probabilities for each genotype using Eqs. 2–5. We do that for each target SNP. We basically create a set of lookup tables with the genotype probabilities (one for each target SNP), where each possible combination of selected tag SNP is an index of the table, and the probabilities of 0 and 1 are the values (the probability of 2 can be calculated from the other probabilities) (Figure 3).

**Figure 3:**
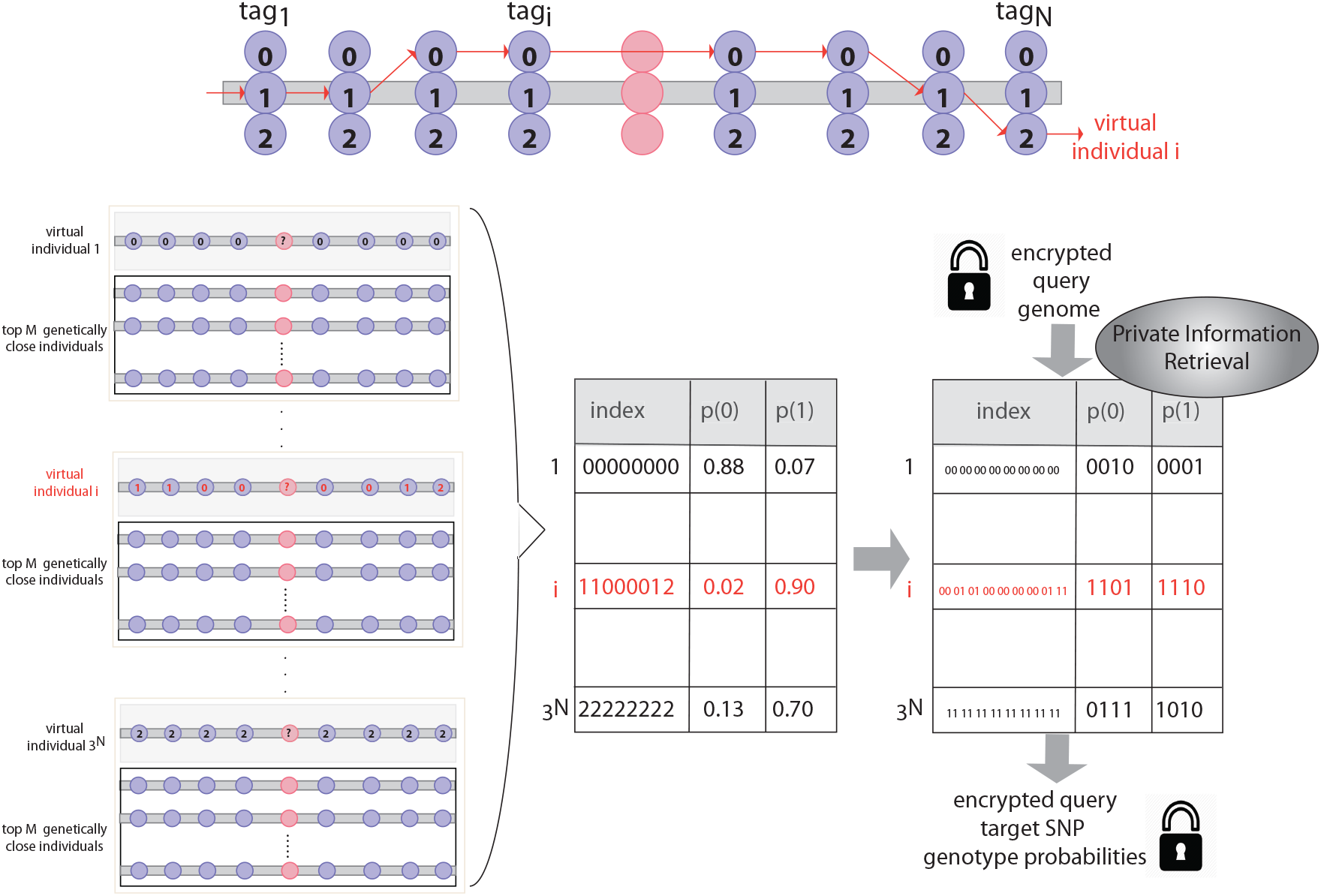
p-Impute algorithm in the encrypted domain. 3^*N*^ *virtual individuals* are generated by using the all possible combination of genotypes of *N* tag SNPs. For each *virtual individual* we find the *M* genetically similar individuals from the training database. We then generate a lookup table, where each index is the sequence of the tag SNP genotypes of a *virtual individual* and the probability of genotypes 0 and 1 for the target SNPs are the columns. These probabilities are calculated using the scoring system described in Figure 2. After converting the look-up table with transformation to the indices and probabilities, we perform *Private Information Retrieval* to the look-up table with encrypted query tag SNP genotypes and return encrypted query target SNP genotype probabilities.

#### Encryption

For each target SNP, we select from each query individual only the tag SNPs relevant for that target SNP (i.e., the *N* closest ones). Let *T* be the number of target SNPs. Then, the total number of tag SNPs per individual to be encrypted is given by *N* · *T*, which is much less than the number of tag SNPs available. Furthermore, each tag SNP contains a value in the set {0, 1, 2}, which we break into its bit representation {00, 01, 10} to allow encrypted comparison during querying. Thus, we have 2*NT* plaintexts per query individual.

Our solution requires addition, multiplication, and comparison on encrypted data. The BFV encryption scheme supports homomorphic addition, subtraction, and multiplication, while comparison can be implemented using homomorphic logic gates, which in turn, can be implemented using additions, subtractions, and multiplications. The implementation of BFV available in SEAL does not support bootstrapping. However, it is not required since we know a priori the number of homomorphic operations executed by our solution. The BFV scheme also supports batching, which allows us to pack many plaintexts in one ciphertext and operate on all plaintexts at the same time, in a single-instruction multiple-data (SIMD) fashion. The number of slots available in a ciphertext is given by the degree of its polynomial modulus, a number in the thousands (see Sections 2.6 and 4 for details on polynomial modulus). We maximize the usage of those slots by packing multiple query individuals into a ciphertext. We pack only independent information in the same ciphertext, i.e., each ciphertext contains only one bit of information related to each target SNP per individual. For example, if we have 500 target SNPs and 2^14^ slots, we can pack 32 individuals in the same ciphertext. And since each target SNP requires *N* tag SNPs and the tag SNPs are broken into two bits each, we need 2*N* ciphertexts to represent those 32 individuals. The number of individuals packed in a ciphertext is given by 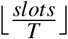.

#### Query

Since the result of our training is a set of lookup tables, our querying consist of indexing these tables. Searching on a table where the query is encrypted is known as the *Private Information Retrieval* problem [19]. In the encrypted domain, the complexity of the search is linear to the number of elements in the table (*O*(*n*)), while exponential in the number of tag SNPs (3^*N*^ is the number of elements in the look-up table, *N* is the number of tag SNPs selected for each target SNP). Despite having one lookup table per target SNP, we can search on all of them in parallel due to batching. Furthermore, all target SNPs of several query individuals are searched in parallel, since they are packed in the same ciphertext in accordance with the encryption methodology.

The first step is to prepare the training data for the querying. We scale the probabilities by a multiplying factor and convert them to integer, and then pack them together for batching. We represent both probabilities (of genotypes 0 and 1) in a single plaintext. We do that by using *shift left* and *add* (e.g. *P*_1,0_ = (*P*(1) ≪ *s*) + *P*(0), where *s* is the shifting constant). Thus, we get all probabilities in a single search, improving performance and reducing memory usage. In our implementation, we use *s* = 7.

In order to perform comparisons in the encrypted domain, we need to convert the data to binary representation, so we can emulate gate operations on top of homomorphic addition, subtraction, and multiplication. That is why we break the genotypes of the tag SNPs into their binary constituents (i.e 0 becomes 00, 1 becomes 01, and 2 becomes 11). We can implement equality using the 2*N* XNOR gates (*x XNOR y* = 1 + 2*xy* − (*x* + *y*)) between the query individual and the table index, and then apply 2*N* − 1 AND gates (*x AND y* = *xy*) to get a binary output. The result of the equality is multiplied by the combined probabilities. If the equality results in the encryption of 1, the result of the multiplication is the probabilities. Otherwise, it is the encryption of zero. Each comparison of the query to a table index will either result in the probabilities for that index or zero. Since it is a lookup table of all possible selected tag SNPs, there is always one and only one match. Therefore, the addition of all partial results is the value of the indexed query. In addition, since AND gate and multiplication are the same operation, we embedded the multiplication between the result of the equality and the value of the table index inside the equality to reduce the depth of the circuit, since deeper circuits produce more noise. In addition, we add thread-level parallelism to the query, where we simply divide the workload (searches) among different threads.

#### Decryption

To get the probabilities and resulting target SNP genotypes, we decrypt the ciphertexts, unpack the data, decouple the probabilities (*shift right* and *mask*), convert them back to floating-point and scale back using the multiplying factor we used in encryption. Then, we calculate the probability for genotype 2 (*P*(2) = 1 − *P*(1) − *P*(0)). The target SNP genotype is the genotype with the highest probability.

The overall procedure for a single target SNP is depicted in Figure 3.

### 2.5 Performance

Our algorithm has two parameters: the number of neighboring SNPs (*N*), and number of genetically similar individuals (*M*). We performed tests with different values of *N* and *M* and evaluated them in terms of accuracy, memory and run time. We then compared the accuracy of the most optimum parameters against the accuracy of the state-of-the-art plaintext methods IMPUTE2 [15] and Beagle [16]. Note that *M* is only used during training and it does not affect the efficiency of our algorithm.

#### Accuracy, memory and running time

For a comprehensive evaluation of the accuracy of the imputation model, we used a metric that reflects both correct predictions (true positive rate) and false mis-classifications (false positive rate) for each of the three genotypes, as was suggested by the iDASH Secure Genome Analysis Challenge. Since this is a prediction problem with three classes, we used Micro-AUC score, which has been used for comparing the accuracy of various multi-class classification/prediction models [20]. We first fixed *N* at 2 to find the best number of individuals *M*. We found that our model has the highest Micro-AUC when *M* is 60 individuals (Figure 4a). We then fixed the *M* at 60 and increased *N* from 2 to 8 and measured not only Micro-AUC but also total roundtrip (encryption+query+decryption) time and memory usage in Figure 4b-d (see Supplementary Figure 1 for time and memory breakdown for encryption, query and decryption separately). We found that Micro-AUC increases only by 0.001 while the time and memory overhead increases exponentially when we increase *N* from 7 to 8. For all three performance measures, *N* equals to 7 seems to be the most optimum parameters in our tests. Since parameter *M* was only used during training, we also showed that the run time and memory performance of query does not depend on *M* (Figure 4). All tests are done using an 8 core 2.10 GHz Intel Xeon processor, using the 9745 tag and 500 target SNPs of Human Chromosome 1.

**Figure 4:**
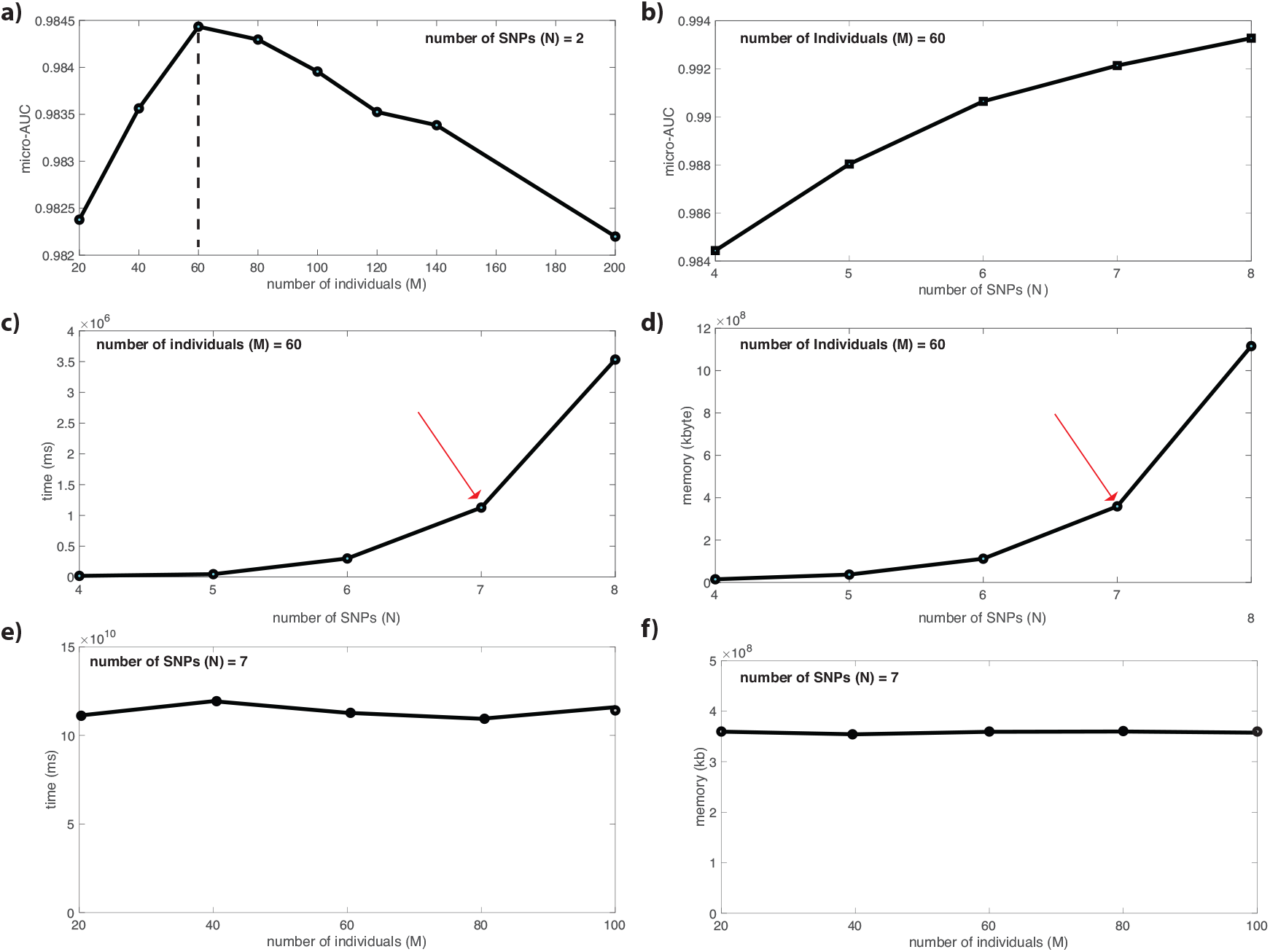
Fine-tuning *N* and *M*: All of the calculations in this figure have been done using the 9745 tag and 500 target SNPs of Human Chromosome 1. **(a)** The change in Micro-AUC with the increasing number of *M*, while *N* is fixed at 2. Micro-AUC reaches a maximum at 60 individuals. **(b)** The change in Micro-AUC with the increasing number of *N*, while *M* is fixed at 60. Micro-AUC increases with the increasing number of neighboring SNPs (*N*). **(c)** The increase in total roundtrip time (encryption+query+decryption) with the increasing number of neighboring SNPs when we fix number of individuals at 60. **(d)** The increase in memory usage with the increasing number of neighboring SNPs when we fix number of individuals at 60. **(e)** The total roundtrip time (encryption+query+decryption) with changing M when we fix the number of SNPs at 7. Since M is only used during training, as can be seen from the plots, it does not affect the runtime. **(f)** The total memory usage with changing M when we fix the number of SNPs at 7. Since M is only used during training, as can be seen from the plots, it does not affect the memory.

#### Comparison to IMPUTE2 and Beagle

We further compared our prediction to the results obtained from running commonly used genotype imputation softwares IMPUTE2 [15] and Beagle [16]. We trained our method, IMPUTE2 and Beagle using the genomes from all 2504 individuals of 1000 genomes project [13] and tested on genomes from 870 individuals of GTEx project [14] by plotting the Receiver Operating Curves (ROC) and calculating the Micro-AUC. We found that the performance of p-Impute is in excellent agreement with the performance of the state-of-the-art plaintext methods (Figure 5). All three algorithms result in a micro-AUC score of 0.99. We also performed the above accuracy test when we have only 1045 tag SNP genotypes to predict the 500 target SNP genotypes of 870 GTEx individuals and found that both p-Impute and IMPUTE2 achieve a Micro-AUC score of 0.97 and Beagle achieves a Micro-AUC score of 0.99. An advantage of our model over Beagle and IMPUTE2 is that we can obtain fast imputation without the need of phasing the genomes, which can be time and compute intensive. Our algorithm in plaintext can be implemented using a hash table, which has complexity *O*(1) per query, i.e providing near instant answer.

**Figure 5:**
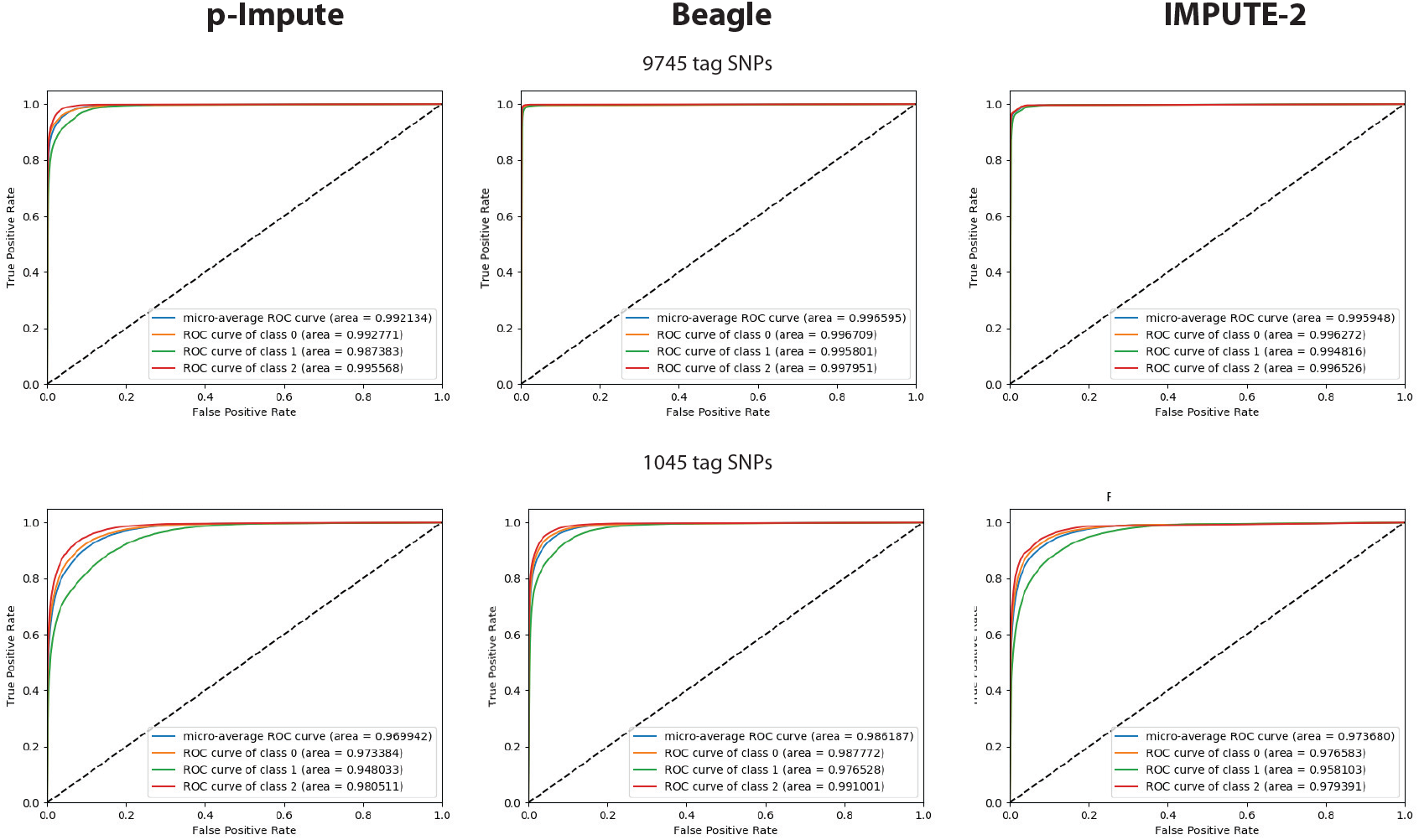
ROCs and Micro-AUC for p-Impute (*N* = 7; *M* = 60), Beagle and IMPUTE2 using GTEx data from Chromosome 1 of 870 individuals. The Micro-AUC has been calculated for predicting genotypes of 500 target SNPs using both 9745 and 1045 tag SNP genotypes.

### 2.6 Security

The security of our implementation depends only on the security of the BFV encryption scheme, which is based on the Ring Learning With Errors (RLWE) problem [21]. We select encryption parameters that provide enough noise budget for the computation and give us 128 bits of security, which is considered secure by the Homomorphic Encryption Security Standard [22]. The plaintext modulus is set to 65537 in order to enable batching. The degree of the polynomial modulus *n* and the coefficient modulus size log_2_(*q*) varies according to the number of select tag SNPs *N*. For *N <* = 4, we use *n* = 2^14^ and log_2_(*q*) = 438 bits, while for *N* > 4, we use *n* = 2^15^ and log_2_(*q*) = 881 bits. See Methods for more details on polynomial modulus.

### 2.7 Code availability

A C++ implementation of our algorithm, the training data as well as the python scripts necessary for performance calculation can be found at https://github.com/gersteinlab/idash19he.

## 3 Discussion

Privacy of individuals’ genomic data has recently emerged as one of the major foci of data privacy studies, as unrestricted availability of personal genetic information gives rise to many concerns. For example, knowledge of genetic predisposition to diseases may bias insurance companies or create unlawful discrimination by employers. Recently it has been also shown that not only DNA sequencing of individuals but also high throughput molecular phenotype datasets, such as functional genomic and metabolomics measurements or even microbiome measurements, increase the number of quasi-identifiers for participating individuals that can be used by adversaries for re-identification purposes. The rise in popularity of genetic and ancestry testing companies, which collect and distribute large amounts of genomics and health data, will only increase privacy concerns over sharing personal biological data.

Human genetics studies mainly focus on identifying genetic variants that affect human traits and diseases. This identification is possible by collecting and analyzing genetics data from large cohorts of individuals with different phenotypes. Genome-wide association studies and studies focusing on identification of various quantitative trait loci (e.g GTEx, psychENCODE) a im to genotype thousands of individuals for a better characterization of human diseases. There are two cost associated bottlenecks for large-scale genomics studies. The first is regarding the feasibility of comprehensively genotyping entire genomes. Although the cost of genome sequencing is rapidly decreasing, it is still not feasible to perform high coverage whole genome sequencing (WGS) on thousands of genomes. To overcome this, researchers developed genotype imputation tools that can predict the missing genotypes using available population genetics data. Although genotype imputation is an extremely powerful technique, it could cause the second bottleneck as it is computationally not feasible to store and impute thousands of genomes in local servers and computers. Therefore, institutions and funding agencies are increasingly outsourcing cloud services for complex genomics analysis such as genotype imputation.

Outsourcing to third party computing environments such as cloud services for storage and analysis of thousands of individuals’ genetic data raises serious privacy concerns. There is an increasing need for fast and scalable privacy-preserving genomic analysis tools. We need tools and software that enable keeping the genetic data encrypted in third party computing environment. This can be achieved by a special type of encryption, called Homomorphic Encryption (HE). HE allows manipulation of encrypted data, hence removing the privacy risk associated with the decryption of sensitive data for computation purposes. However, there is a significant computational overhead incurred in HE calculations compared to computations on plaintext. This cost depends on the class of computation needed on the encrypted variables. When arbitrary computation is required (which is the case for most of genomic calculations), Fully Homomorphic Encryption (FHE) needs to be employed, but at tremendous overheads (several orders of magnitude slowdown). Therefore, it is important to design fast and scalable algorithms that can be implemented using FHE.

In this study, we developed a scalable privacy-preserving genotype imputation method using FHE scheme. We showed that our imputation results are in excellent agreement with its plaintext counterparts when tested on 870 genomes. Since Chromosome 1 is one of the largest chromosomes in the genomes and each chromosome can be imputed independently and in parallel, we showed that we can achieve genome-wide imputation in under 1.5 days in the encrypted domain for 870 individuals.

## 4 Methods

### 4.1 Homomorphic Encryption

HE allows the possibility of the computing on encrypted data directly without decryption. There are variety of mathematical models for HE, that support different kinds and number of operations. Some of these models support one operation such as the Paillier encryption scheme [23] of partial homomorphic encryption. The scheme supporting more than one operation allow theoretically all possible computation of recursive functions, and is called Fully Homomorphic Encryption (FHE) scheme.

The security of fully homomorphic encryption is provided by the hardness of the RLWE problem. The hardness of the RLWE problem is due to adding a small amount of error to a point in a lattice, which makes it difficult to determine which point that error was added to. This creates noise, hence the ciphertexts in homomorphic encryption schemes are noisy, which grows during homomorphic additions and multiplications. This growth eventually makes it impossible to decrypt the resulting ciphertext in FHE. For that, Gentry proposed a bootstrapping technique which can evaluate its own decryption function [24]. Bootstrapping re-creates a ciphertext by running the decryption function on it homomorphically with encrypted secret key, which reduces the noise.

Although bootstrapping is extremely helpful in deep arithmetic circuits, if the number of operations needed in an algorithm is known and small, bootstrapping may not be needed. Some of the leveled FHE constructions do not use bootstrapping procedure. A leveled FHE scheme can evaluate *L*-level arithmetic circuits with *O*(*λ* · *L*^3^) per-gate computation. Security is based on RLWE for an approximation factor exponential in *L*.

Data in these encryption schemes is represented by polynomials both when it is encrypted (the ciphertext) and when it is unencrypted (the plaintext). A special polynomial called the polynomial modulus is defined, and only the remainder of polynomials is considered when they have been divided by this polynomial modulus.

### 4.2 Brakerski/Fan-Vercauteren (BFV) encryption system

The BFV encryption system [11] is a lattice-based cryptographic scheme that depends on the hardness of the RLWE problem. With BFV, one can perform both addition/subtraction and multiplication within the encrypted domain. Divisions and exponentiation of a number by an encrypted one and non-polynomial operations are not supported. The computations can only be performed on integers. Eq. 6 depicts the concept of homomorphism, where *c*_1_ and *c*_2_ are ciphertexts, *E* and *D* are encryption and decryption functions, respectively, and ⊗ and ∗ are homomorphically equivalent operators, i.e., ⊗ over ciphertexts is equivalent to the encryption of ∗ over plaintexts.

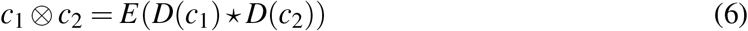

Bootstrapping is supported by BFV, but due to its inefficiency, it is not commonly implemented by FHE libraries such as Microsoft SEAL, making it in practice a Somewhat Homomorphic Encryption scheme. This limitation leads to the need of knowing the depth of the arithmetic circuit for properly selecting the encryption parameters.

The specific form of this polynomial modulus in the BFV scheme is *x*^*d*^ + 1. where *d* = 2*n* for some *n*.

